# A Pilot Study to Establish a Penetrating Traumatic Brain Injury Rat Model for Implantation of a 3D Printed Scaffold

**DOI:** 10.1101/2025.03.20.644358

**Authors:** Meaghan E. Harley-Troxell, Michelle Dennis, Madhu S. Dhar

## Abstract

**Introduction:** Traumatic brain injuries (TBIs) are the leading cause of death and disability, with penetrating TBIs being the most lethal form. As the primary injury involves a foreign object breaking the skull, disrupting the blood brain barrier (BBB), and damaging the brain tissue, the secondary injury that follows is further damaging with persistent inflammation leading to tissue atrophy. While no TBI treatments currently exist, ongoing investigations are developing biomaterial scaffolds and cellular therapies to improve upon the poor outcomes from this disease. This pilot study sets out to establish a TBI rat model that maintains focal damage to the cerebral cortex, while manually disrupting the BBB. Injuries disrupting this barrier need to be managed differently than those that do not, allowing us to develop a specific, therapeutic treatment for this type of injury. We hypothesize that our method of BBB disruption will indicate behavioral, physical, and histological evidence of a TBI. Our TBI model will also create a cranial opening in which we can ensure surgical feasibility of implantation of a scaffold. We hypothesize the implantation of FDA-approved synthetic polymer, poly (lactic-co-glycolic acid) (PLGA), and carbon-based nanomaterial, reduced graphene oxide (rGO), will not show evidence of a foreign body rejection at 30 days after surgery.

**Methods:** Four Sprague Dawley rats underwent a stereotaxic surgery with a 5-mm craniotomy. The dura and brain tissue were disrupted using a beaver blade. The PLGA/rGO scaffold was gently placed onto the brain tissue. Neurological function was evaluated for the first three days, then weekly throughout the 30-day study. At 30 days, brains were dissected, paraffin embedded, and sectioned for H&E and Prussian blue staining, and immunohistochemistry (IHC).

**Results:** Neurological function assessments indicated no change in rat behavior and normal wound healing over the 30 day study. H&E and Prussian blue staining indicated mild leptomeningeal thickening and evidence of hemosiderin in 3 rats. One rat had foreign body giant cells and an abscess around the implanted material with evidence of more severe leptomeningeal thickening and hemosiderin. IHC indicated normal anatomic structures with no changes in 5 of the 6 markers at 30 days after surgery. Neural marker, NeuN, had a significant decrease in expression for all four rats.

**Discussion:** While there was no behavioral or symptomatic evidence of a TBI, histology showed evidence of a mild, focal TBI in 3 of the 4 rats, and evidence of a foreign body response and a severe, focal TBI in 1 rat. Future studies will perform IHC at earlier timepoints to confirm additional biomarkers, and will implant a scaffold that is more mechanically aligned with the brain tissue to further evaluate the biocompatibility of graphene nanoparticles in brain tissue, and the effectiveness of a therapeutic scaffold.

## 1. Introduction

Traumatic brain injuries (TBIs) are the leading cause of death and disability (1, 2). Annually in the United States, approximately 1.7 million people have a TBI event, resulting in 50,000 deaths (1). Currently it is estimated that 3.2 – 5.3 million are currently living with a TBI-related disability (1). Penetrating TBIs are the most lethal form, with a foreign object breaking through the skull, disrupting the blood brain barrier (BBB) and damaging the brain tissue (3, 4). These are most often caused by accidents, including household firearms, falls, and motor vehicle collisions (2-4). The severity of the primary injury is worsened by circulating immune cells foreign to the local environment crossing the disrupted BBB, causing persistent inflammation and generation of reactive oxygen species (ROS) that result in loss of neural tissue and damaged vasculature (1, 2, 5). This damage results in severe clinical symptoms including high cranial pressures and hemorrhage. While no treatments for TBI exist, management strategies include various neurosurgical interventions, and pharmacological and non-pharmacological methods (2-4, 6-8). While surgical interventions may improve survival outcomes for severe TBIs, they come with increased risk of infection, longer recovery times, and may not result in greater functional outcomes (3). Currently researchers are investigating tissue engineering and regenerative medicine strategies to develop biomaterial scaffolds and cellular therapies that may improve upon these poor outcomes (2, 6, 7, 9).

Animals are a necessary preclinical step for translational research in the development of novel therapeutics, and can generate TBI solutions for both animal and human medicine (10). While large animals, such as pigs, are important models for TBI, showing more complexity, and grey and white matter ratios similar to human brains; using rodents, such as rats or mice, prior to large animal models are a crucial step (11, 12). Outside the advantages of low costs and easy handling, rats offer a high level of standardization, and can be assessed for functional outcomes and the pathophysiology of cellular damage, in a way that mimics the human brain (11). Several preclinical models exist to mimic TBIs including controlled cortical impact injury models, weight drop models, and penetrating ballistic-like models (13). There are a wide range of TBIs, and while each model excels in representing a specific type of scenario, they do not consistently disrupt the BBB, while maintaining controlled, focal damage to the cerebral cortex as intended in this study (13, 14). Severe mechanical models often impact the brain as a whole, simulating concussion-like injuries alongside the penetrating impact, and resulting in high levels of fatality (14). Meanwhile, the penetrating ballistic-like model, while remaining focal, creates a deep cavity mimicking a combat setting. Adding to the available preclinical models to accurately represent the range of TBIs may alter how clinical care is managed, and be useful in evaluating more specific, novel therapeutics (15).

In this preliminary study we performed a craniotomy in rats to manually cut the BBB, managing the disruption to the brain tissue. This allowed us to assess the feasibility of our therapeutic scaffold implantation, and safety of the biomaterial-tissue interaction between the brain tissue and our proposed scaffold containing poly (lactic-co-glycolic acid) (PLGA) and graphene. The use of stereotaxic surgical equipment and cranial landmarks, allowed for a standardized injury, capable of replication. We used a neurological assessment and immunohistochemistry to identify if disruption of the BBB was sufficient to model a TBI, while identifying if a foreign body reaction occurs with the implanted materials. While PLGA is an FDA-approved synthetic polymer, graphene’s biocompatibility is highly variable, depending on a variety of tunable physicochemical and mechanical properties (16). For this study, we used a previously fabricated scaffold to evaluate the graphene’s biocompatibility with the brain tissue. We hypothesized that the disruption of the BBB would cause behavioral, physical, and histological evidence of a TBI, while the implanted material would not show evidence of a foreign body rejection or chronic inflammation 30 days after surgery.

## 2. Methods

### 2.1. Biochemicals, Chemicals, and Disposables

All biochemicals, cell culture supplements, and disposable tissue culture supplies were purchased from Thermo Fisher Scientific (Waltham, MA, USA) unless otherwise noted.

### 2.2. Animals

Sprague Dawley rats (Charles River Laboratories, n=4, male, aged 6 months) were acclimated prior to beginning procedures. All procedures were conducted in accordance with PHS guidelines for the humane treatment of animals under approved protocols established through the University of Tennessee’s Institutional Animal Care and Use Committee (IACUC; protocol # 2954-0623).

### 2.3. Surgical Procedure

Rats were anesthetized using inhalant isoflurane, and buprenorphine (0.05 mg/kg) was administered subcutaneously prior to surgery. The surgical area was shaved, and eye lubricant was applied. Under aseptic conditions, the rat was mounted onto the stereotaxic apparatus (KOPF, Tujunga, CA, USA), using a nose cone to maintain anesthesia throughout the procedure (7, 17, 18). The dorsal/ventral adaptor was aligned to the -8 setting; the ear bars were aligned to the 7.5 setting on either side. The area was draped, and the scalp was cleaned with iodine prior to cutting a 1-2 cm longitudinal incision to expose the skull. The periosteum was gently teased back from the skull and the area was cleaned with 95% ethanol. A handheld micro-drill (Stoelting Co., Wood Dale, IL, USA) with a sterile 5-mm drill bit was used to create a craniotomy to the left of the sagittal suture, midway between the lambda and bregma landmarks. The dura matter was damaged by cutting the layer using forceps and a beaver blade. A poly (lactic-co-glycolic acid) (65:35 PLGA) (Sigma-Aldrich, St. Louis, MO, USA) and 0.5% reduced graphene oxide (rGO) (Cheap Tubes Inc., Grafton, VT, USA) scaffold from a previous study was cut to the 5mm size and implanted on the cerebral cortex (19). Briefly, PLGA and rGO were blended with 0.5 mL dimethyl sulfoxide (DMSO), and melted into a homogenous mixture for extrusion-based 3D-printing using the Cellink-BIO X6™ printer. The scaffold was printed in 15 filament layers, with approximately 80% porosity, and printed with 5 mm (x), 5 mm (y), and 2 mm (z) dimensions. After implantation, the area was covered with bone wax and the scalp was closed with 4-0 absorbable surgical sutures (Ethicon Inc., Raritan, NJ, USA). Rats recovered on a heating pad, were individually housed, and monitored twice daily for the first 3-days, daily for 1-week, and weekly throughout the remaining 30-day study. Buprenorphine was administered every 12-hours for 3-days after surgery, and ketoprofen (5 mg/kg) was administered every 24-hours for 3-days after surgery. Baytril (100 mg/400 mL) mixed into water with flavored Gatorade, was provided for 1-week after surgery.

### 2.4. Neurological Function Evaluation

Animal recovery and neurological function were evaluated for each rat on days 1, 2, 3, 7, 14, and 21 after surgery. A modified neurological severity score was used to assess the body weight, general condition, physical appearance, breathing, spontaneous behavior, handling reaction, wound healing, and neurological evaluation of each rat (11, 20-22). These behaviors were graded on a scale of 0 (normal) to 30+ (severe) to determine any neurological deficits that occurred from the cranial intervention or material implantation.

### 2.5. Immunohistochemistry of Brain Tissue

At the 1-month end-of-study timepoint, the intact brain was removed as previously described and placed in 10% formalin (23, 24). The cerebellum was removed, and the brain was dissected down the midsagittal plane, dividing the left and right hemispheres of the cerebral cortex. Each half was paraffin embedded, and longitudinally sectioned. Sections were deparaffinized, hydrated, unmasked, stained, and mounted as previously described (16). One histology section from all specimens were stained with hematoxylin and eosin (H&E) (Azer Scientific, Inc., Morgantown, PA, USA) for assessment of cellular detail. One histology section from all specimens were stained with Prussian Blue for assessment of hemosiderin deposition, using a 30 minute stain with 1:1 hydrochloric acid (HCl) and potassium ferrocyanide (KCn). Immunohistochemistry (IHC) was performed on remaining sections using: glial fibrillary acidic protein (GFAP) (1:500), neuronal nuclei (NeuN) (1:500; Cell Signaling Technology, Inc., Danvers, MA, USA), von Willebrand factor (vWF) (1:1000; Abcam, Cambridge, UK), neurofilament light chain (NEFL) (1:20), ionized calcium-binding adapter molecule (Iba1) (1:500), and CD68 (1:500). The respective secondary antibodies used were: goat anti-rabbit IgG HRP (1:500), and rabbit anti-mouse IgG HRP (1:500; Abcam, Cambridge, UK). Briefly, GFAP identifies astrocytes, NeuN identifies neurons, vWF identifies vascularization, NEFL identifies neural cytoskeleton structure, Iba1 identifies microglia, and CD68 identifies phagocytic activity (25-30). Sections were imaged using Keyence BZ-X Series All-In-One Fluorescent Microscope (Keyence Corporation of America, Itasca, IL, USA) at 5X, 10X, 20X, or 40X magnification. The H&E sections were analyzed by a trained histologist. For each section, multiple images were taken along the length of the section with 20% overlap and stitched together to create a single image of the entire section. Scale bars were added. ImageJ software was used to analyze the stitched images (31). They were selected for optimal contrast (blue) and threshold (range: 150-250). The nerve was isolated from the background and converted to a black and white image (black = stained tissue). For each section the nerve was measured for percent area (percentage of black in the image; %) (16). Mean and standard deviation values were obtained. GraphPad Prism (version 10) was used to perform a Shapiro-Wilk test of normality prior to t-tests for statistical significance (p ≤ 0.05).

## 3. Results and Discussion

Carbon nanomaterial, rGO, has been investigated as a component for nerve tissue engineering due to its rough surface area distributed with oxygen-containing functional groups that influence its tunable mechanical properties, and positive influence on cell growth (32, 33). rGO can be engineered using a variety of 3D printing techniques to form scaffolds that attach, proliferate, and differentiate viable exogenous or endogenous stem and progenitor cells to aid in nerve repair and regeneration (33-35). Particularly, rGO features the unique property of electrical conductivity to improve functional nerve restoration (32, 33, 36). While many studies have focused on graphene’s potential in the peripheral nervous system, graphene and its derivatives have also shown to be a promising component for central nerve injuries (37-39). Controversy surrounding rGO’s biocompatibility has slowed its translation to clinical use. Numerous variables such as surface functionalization, particle shape and size, dispersion, concentration, dosage, route of administration, and processing techniques can all influence how cells respond to rGO (32, 36, 39- 45). Therefore, we must evaluate if our specific rGO-containing construct is safe to use in this delicate tissue model.

A TBI is a complex disease in which a primary injury results in an immediate disruption of brain tissue caused by an external source, leading to contusion, hemorrhage, and axonal shearing. The following secondary injury evolves over the following hours, days, and months, involving a cascade of degenerative molecular events, leading to cell death (13, 35). Patients who experience focal, penetrating TBIs have areas of brain laceration, disruption of the leptomeninges, subdural hemorrhage, and cerebral edema (29). Our method of injury manually disrupted the dura mater and cerebral cortex with a small blade through the cranial opening (**Figure 1A**). When the scaffold was placed onto the exposed brain tissue for rats 1, 2, and 4, there was minor bleeding before the scaffold was placed against the brain tissue with mild pressure. With rat 3, there was heavy bleeding after the surgical procedure, and the scaffold was placed onto the brain tissue after ensuring the bleeding was under control. Evidence of the differences in the surgical implantation were seen 30 days later during the brain dissection. In rats 1, 2, and 4 the scaffold was more strongly incorporated into the surrounding bone of the craniotomy than with the brain tissue (**Figure 1B**). Some theories as to why this may have occurred include the location of the implanted material, which may not have been pushed sufficiently through the cranial opening, the large z axis of the implanted scaffold may have interfered with the implantation process, or excessive cerebral pressure or edema may have pushed the construct back into the opening due to insufficient closing using the bone wax. Meanwhile rat 3 showed a greater incorporation into the brain tissue (**Figure 1C**). This may be due to the more severe damage to the brain, evident with the heavy bleeding and deeper implantation of the scaffold.

**Figure 1.**
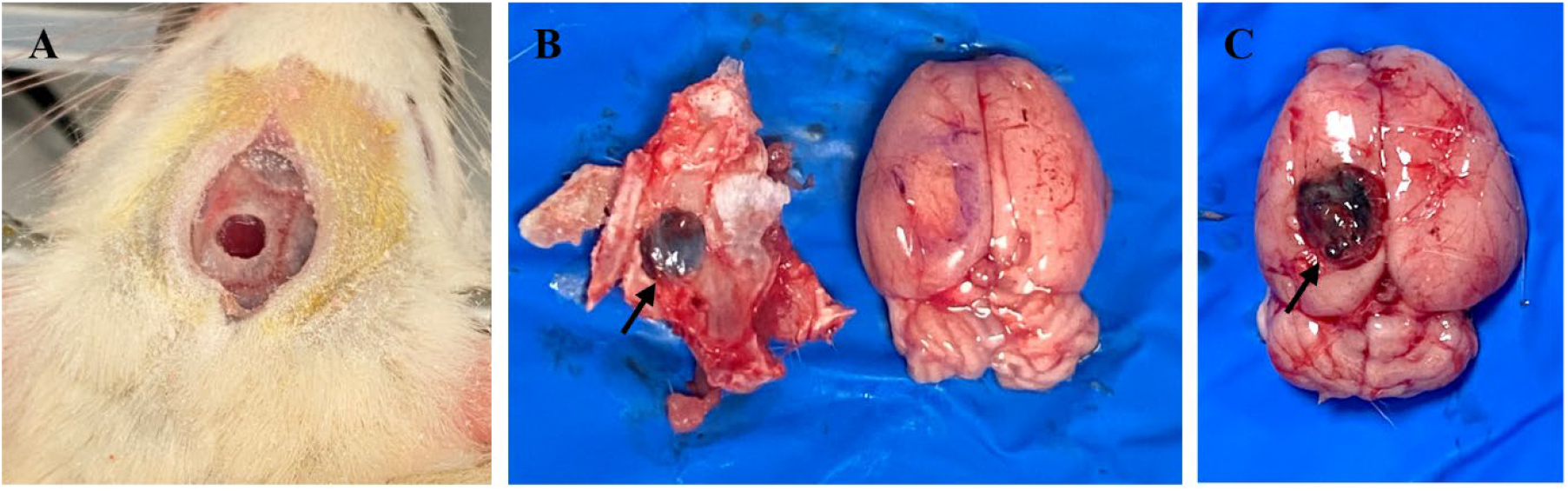
**A**. Image of the craniotomy location. **B**. Brain dissection of rat 4; tissue pen used to circle area of injury; black arrow indicates implanted material. **C**. Brain dissection of rat 3; visualizes deeper cortex injury and material implantation; black arrow indicates implanted material.

During the 30 day study, the neurological assessment evaluated the physical and behavioral changes that may occur with a TBI (11, 20-22). At all six time points, all rats showed no changes in any of the areas assessed (**Table 1**). Rat 3 took slightly longer to come out from anesthesia, which was expected due to the longer procedure time to control the heavy bleeding. However, while rat 3 showed slightly slower spontaneous movement, it still fell within normal parameters.

**Table 1.** Neurological evaluation score sheet indicates response for all rats at all time points throughout the 30 day study. Evaluation indicates no abnormal behavior or physical reactions after surgery, injury, and material implantation.

| Neurological Evaluation Score Sheet |  |  |  |  |  |
| --- | --- | --- | --- | --- | --- |
| Point of Evaluation | Score Scale | Rat 1 | Rat 2 | Rat 3 | Rat 4 |
| Loss of Body Weight | <5% (0 points) | X | X | X | X |
|  | 5-10% (1 points) |  |  |  |  |
|  | 11-15% (5 points) |  |  |  |  |
|  | 16-20% (10 points) |  |  |  |  |
|  | >20% (30 points) |  |  |  |  |
| General Condition, Physical Appearance | Well groomed, clean fur, clean and intact wound, clear eyes (0 points) | X | X | X | X |
|  | Coat slightly unkempt, eye redness/ “red tears” (chromodacryorrhea) (1 points) |  |  |  |  |
|  | Coat dirty and shaggy, orbital tightening, nose flattening, ear and whisker changes, skin-pinch (dehydration test), some goosebumps (piloerection) (5 points) |  |  |  |  |
|  | Coat unkempt, piloerection (10 points) |  |  |  |  |
| Breathing | Normal breathing (0 points) | X | X | X | X |
|  | Abnormally rapid breathing (tachypnea) (5 points) |  |  |  |  |
|  | Difficult or labored breathing (dyspnea) (10 points) |  |  |  |  |
| Spontaneous Behavior | Normal movements/locomotion (0 points) | X | X | X | X |
|  | Reluctance to move, slightly abnormal gait (5 points) |  |  |  |  |
|  | Lethargic, apathetic, abnormal gait (10 points) |  |  |  |  |
|  | Significant mobility problems, mobility intermittent <12 hours (20 points) |  |  |  |  |
|  | Immobile >12 hours (30 points) |  |  |  |  |
| Handling Reaction | Normal curiosity, alert (0 points) | X | X | X | X |
|  | Tense and nervous when held (5 points) |  |  |  |  |
|  | Markedly distressed when handled (e.g., shaking, vocalizing, aggression) (10 points) |  |  |  |  |
| **Wound Healing** | Wound clear, sutures/wound clip in place, no signs of infection (0 points) | X | X | X | X |
|  | Wound closure insufficient, open wound, no signs of infection (5 points) |  |  |  |  |
|  | Wound showing initial signs of infection - swelling and redness (10 points) |  |  |  |  |
|  | Wound infection severe – swelling, redness, discharge (25 points) |  |  |  |  |
| **Neurological Evaluation** | No observable deficit (0 points) | X | X | X | X |
|  | Forelimb flexion (paws pulled in when held by tail) (1 points) |  |  |  |  |
|  | Decreased resistance to lateral push (slow/less movement/push-back when side is pushed) + forelimb flexion (5 points) |  |  |  |  |
|  | Circling (circular movement when held by tail) + decreased lateral push and forelimb flexion (10 points) |  |  |  |  |
|  | Seizure, coma, complete paralysis (30 points) |  |  |  |  |
| **Total Score** | Score 0-9: minimal/no action required. | X | X | X | X |
|  | Score 10-20: provide supplementary care as needed. |  |  |  |  |
|  | Score 21-29: consult veterinarian for additional care. |  |  |  |  |
|  | Score 30+: implement animal removal. |  |  |  |  |

While this indicated no immediate concerns regarding a foreign body rejection of the implanted PLGA/rGO scaffold, it also indicated the TBI model may not have been severe enough to implicate neurological deficits common with the disease. Other than TBI severity, the points of neurological evaluation may be expanded to include additional behavior tests, identifying more minute behavioral changes in future studies. Monitoring of intercranial pressure throughout a future study may also aid in confirmation of a TBI, as a signature symptom of the injury.

At the end of the 30 day study, H&E stained sections were evaluated for local tissue reactions. Similar to the lateral fluid percussion injury model where the TBI is focused to one cerebral hemisphere, the contralateral side may be used as a comparison for neural injury (12, 13). Since the injury was focused to the left cerebral cortex, the right cerebral cortex was used as uninjured control tissue. The H&E stained tissue confirmed that no significant pathology was identified in the control tissues for all four rats (**Figure 2A**). For the injured side, rats 1, 2, and 4 showed a slight indentation to the cortex with mildly thickened leptomeninges, reactive small blood vessels, increased stromal cells, and hemosiderin deposition (**Figure 2B,C**). Hemosiderin deposition is a sign of injury, often occurring after hemorrhage, and was confirmed with Prussian blue staining. Together these findings indicate a mild focal cerebral meningeal fibrosis injury. This type of chronic response may occur after a penetrating TBI (46). Rat 3 showed a cavitated region of the cerebrum where the meninges was severely expanded by a mass (**Figure 2D-F**). The mass is composed of neutrophils and necrotic cell debris, marginated by histiocytic infiltrate, and a periphery of fibrovascular tissue with aggregated plasma cells, lymphocytes, and occasional multinucleate foreign body giant cells (FBGCs). Some of the rGO deposits in the area are associated with distinct vacuoles. Hemosiderin deposition was also evident and confirmed with Prussian blue staining. No evidence of infectious agents were present. Overall, rat 3 had evidence of severe focal chronic pyogranulomatous meningitis and a foreign body reaction to the rGO nanoparticles. It was difficult to decipher if the injury was more severe in rat 3, or if the strong inflammatory and immune response were solely due to the scaffold. Future studies will further investigate the separation in this data by using a sham surgical group, and an untreated TBI group.

**Figure 2.**
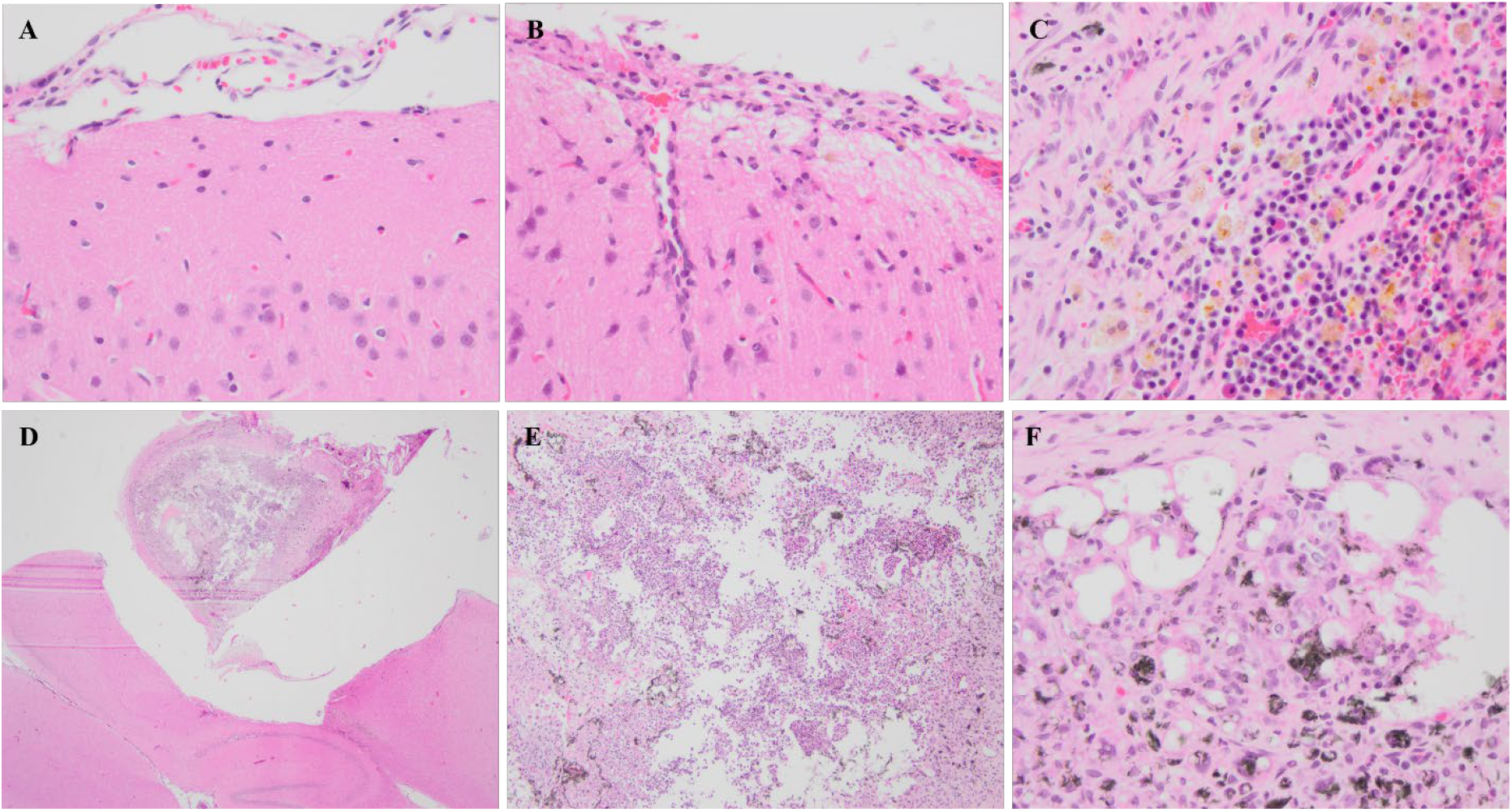
Hemoxylin and Eosin (H&E)-stained sections of brain tissue. **A**. Normal, healthy brain tissue on the control side at 40X magnification. **B**. Mild meningeal thickening and evidence of hemosiderin in rat 2, injured side at 40X magnification. **C**. Brown/orange hemosiderin deposits and black graphene material deposits in rat 3, side A at 40X magnification. **D**. An image at 2X magnification, **E**. an image at 10X magnification, and **F**. an image at 40X magnification exhibiting an abscess surrounding the graphene material implant in rat 3.

The IHC markers that were evaluated have previously shown changes post-TBI. Glial markers, GFAP and Iba1, increase, while neural marker, NeuN, decreases after an injury, due to glial scarring and the loss of damaged neural cells in the area, respectively (26, 27, 29). vWF increases to identify vascular damage after a severe injury, but typically maintains its expression with a mild injury (26). NEFL increases to identify fragmented neural structures, and CD68 increases to indicate a rise in phagocytic activity (25, 29). Qualitatively, each of these six markers showed “normal” anatomical structures (**Figure 3A-F**). Quantitatively, GFAP, vWF, NEFL, Iba1, and CD68 showed no significant differences in expression between the injured and control tissues (**Figure 3G**). However, NeuN expression significantly decreased from 9.65 to 4.56 percent area stained in the injured tissue in all rats (p-value 0.017). A significant decrease in NeuN is expected after a TBI, with the associated loss of neural cells due to the sustained damage to the tissue (26, 29). When rat 3 was separated out from the statistical analysis, the changes in statistical significance were unchanged. The remaining markers may not have indicated any changes due to the 30-day time frame. It has been shown that these markers return to pre-injury levels by 14 days after a TBI in mild injuries (29, 30).

**Figure 3.**
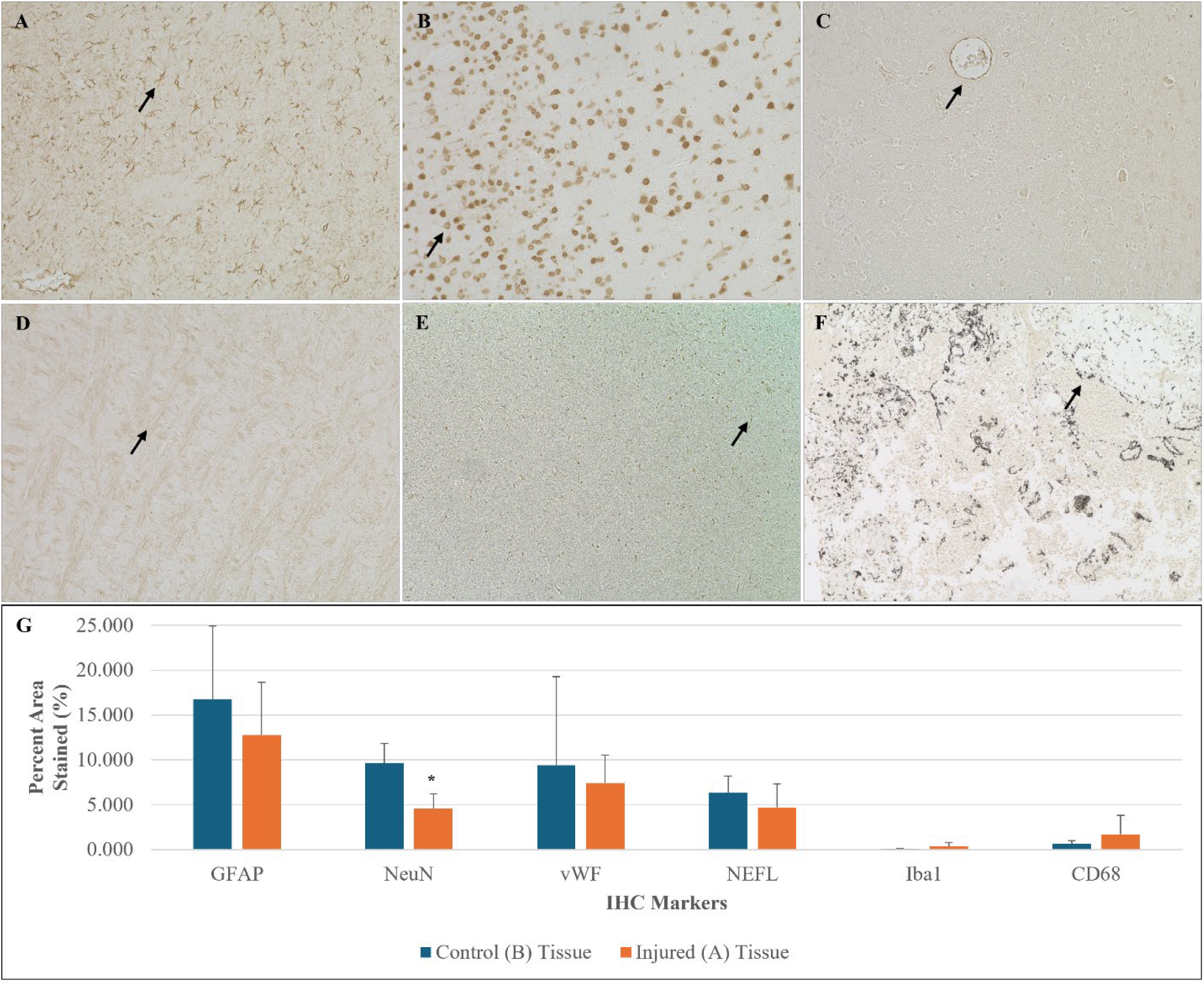
Immunohistochemistry of injured brain tissue. **A**. Representative image of GFAP expression at 20X magnification; black arrow indicates spindle like astrocytes. **B**. Representative image of NeuN expression at 20X magnification; black arrow indicates hollowing of neurons. **C**. Representative image of vWF expression at 20X magnification; black arrow indicates blood vessel. **D**. Representative image of NEFL expression at 20X magnification; black arrow indicates neurofilament alignment throughout tissue. **E**. Representative image of Iba1 expression at 10X magnification; black arrow indicates expression by spindle like microglia. **F**. Representative image of CD68 expression at 10X magnification; black arow indicates phagocytic activity around graphene material. **G**. Quantification of IHC staining for each marker, comparing the control ipsilateral cortex tissue to the injured cortex tissue. No significant changes were seen in GFAP, vWF, NEFL, Iba1, and CD68 expression. NeuN had a significant decrease in expression at 30 days after injury as compared to the control tissue. *p ≤ 0.05.

Overall, all rats showed evidence of a TBI histologically at 30 days after surgery, though they were asymptomatic according to the neurological assessment. Rats 1, 2, and 4 showed evidence of a mild, focal, penetrating TBI according to the H&E staining and IHC marker, NeuN. Rat 3 showed evidence of foreign body rejection that could not be separated from evidence of a severe TBI. While this pilot study did not use a surgical control group, the use of a sham surgery in a larger study, and with a greater number of animals for statistical analysis can separate the data to determine if the injury identified with the H&E staining was due to a severe TBI or the implanted scaffold, and if the variability seen in one of the four rats evaluated is a concern in this model of TBI. Additionally, the surgical procedure was successful for scaffold implantation, though a construct with mechanical properties more aligned with that of the cortex will aid in the successful integration into the appropriate tissue (9, 47, 48). Thirty days was chosen as this is the estimated time a scaffold should remain implanted before degradation (49). This timeframe best allows for the alignment of complete degradation with tissue repair and regeneration in the cerebral cortex. The implanted scaffold did not fully degrade at 30 days in vivo, and will need to be altered to accelerate degradation within this timeframe. Future studies can evaluate the surgical groups at multiple timepoints, including a shorter timeframe, such as 24 hours or 3 days, to better confirm the acute injury (0-3 days) with TBI biomarkers, such as those evaluated with IHC (29, 30, 35). This will also allow for the evaluation of the therapeutic potential of implanted scaffolds and graphene nanoparticles at the acute and chronic TBI phases (9).

## Author Contributions

M.E.H.-T., M.D., and M.S.D were responsible for interpreting the data. M.E.H.-T. and M.S.D. were responsible for conceptualizing the idea and editing the manuscript. M.D. and M.S.D. were responsible for methodology. M.E.H.-T. was responsible for data acquisition and analysis, visualization, and the initial manuscript preparation. M.S.D was responsible for supervision, resources, and writing review. All authors have read and agreed to the published version of the manuscript.

## Data Availability Statement

The datasets generated for this study are available from the corresponding author upon reasonable request.

## Acknowledgments

The authors acknowledge Matthew Cooper for the training on the stereotaxic surgical equipment.

